# Targeted Reactivation of *FMR1* Transcription in Fragile X Syndrome Embryonic Stem Cells

**DOI:** 10.1101/286732

**Authors:** Jill M. Haenfler, Geena Skariah, Caitlin M. Rodriguez, Andre Monteiro da Rocha, Jack M. Parent, Gary D. Smith, Peter K. Todd

**Author notes:** Correspondence: Dr. Peter Todd. The authors wish it to be known that, in their opinion, the first 2 authors should be regarded as joint first authors.

## Abstract

Fragile X Syndrome (FXS) is the most common inherited cause of intellectual disability and autism. It results from expansion of a CGG nucleotide repeat in the 5’ untranslated region of *FMR1.* Large expansions elicit repeat and promoter hyper-methylation, heterochromatin formation, *FMR1* transcriptional silencing, and loss of the Fragile X protein, FMRP. Efforts aimed at correcting the sequelae resultant from FMRP loss have thus far proven insufficient, perhaps because of FMRP’s pleiotropic functions. As the repeats do not disrupt the FMRP coding sequence, reactivation of endogenous FMR1 gene expression could correct the proximal event in FXS pathogenesis. Here we utilize the CRISPR/dCAS9 system to selectively re-activate transcription from the silenced FMR1 locus. Fusion of the transcriptional activator VP192 to dCAS9 robustly enhances FMR1 transcription and increases FMRP levels when targeted directly to the CGG repeat in human cells. Using a previously uncharacterized FXS human embryonic stem cell (hESC) line which acquires transcriptional silencing with serial passaging, we achieved locus-specific transcriptional re-activation of FMR1 mRNA expression despite promoter and repeat methylation. These studies demonstrate that FMR1 mRNA expression can be selectively reactivated in human patient cells, creating a pathway forward for therapeutic development in Fragile X Syndrome.

## INTRODUCTION

Fragile X Syndrome (FXS) is an X-linked disorder affecting approximately 1 in 4,000 males and 1 in 8,000 females worldwide (Tassone, Iong et al. 2012, Hunter, Rivero-Arias et al. 2014). It is the leading inherited cause of intellectual disability and autism. Many FXS patients also experience Attention Deficit Hyperactivity Disorder (ADHD), increased seizure susceptibility, anxiety, and language difficulties. FXS results from expansion of a CGG trinucleotide repeat within the 5’ untranslated region (UTR) of the fragile X gene, *FMR1*. Normally, *FMR1* has between 25 and 40 CGG repeats. Instability of the CGG repeat over multiple generations leads to large (>200) expansions that markedly alter the epigenetic profile of the *FMR1* locus (reviewed in (Usdin and Kumari 2015)). In most FXS patients, both the CGG repeat and the *FMR1* promoter are hypermethylated at cytosine residues (Oberle, Rousseau et al. 1991, Pieretti, Zhang et al. 1991). This hypermethylation is associated with epigenetic marks consistent with heterochromatin formation over the locus and a partial or complete loss of *FMR1* mRNA transcription (Coffee, Zhang et al. 2002). Although the exact mechanism and order of events leading to transcriptional silencing remains incompletely understood, the net result of these epigenetic alterations is the absence of the fragile X mental retardation protein, FMRP.

There is strong evidence that loss of FMRP causes FXS symptoms, as rare patients with mutations or deletions elsewhere in *FMR1* also present with FXS (Gedeon, Baker et al. 1992, De Boulle, Verkerk et al. 1993, Bhakar, Dolen et al. 2012, Santoro, Bray et al. 2012). Moreover, *Fmr1* KO mouse models recapitulate many key features of the human disease, including learning deficits, abnormal socialization and anxiety behaviors, enhanced seizure susceptibility, and dendritic spine morphologic abnormalities(Bhakar, Dolen et al. 2012, Santoro, Bray et al. 2012). FMRP is an RNA-binding protein that binds ~4% of brain messenger RNAs (mRNAs), including an enriched fraction of synaptic transcripts from genes associated with autism (Brown, Jin et al. 2001, Darnell, Van Driesche et al. 2011, Ascano, Mukherjee et al. 2012). FMRP regulates activity-dependent protein translation at synapses (Bhakar, Dolen et al. 2012), where it suppresses translation of bound transcripts, either through direct interactions or via association with translating ribosomes (Feng, Gutekunst et al. 1997, Darnell, Van Driesche et al. 2011, Chen, Sharma et al. 2014). Upon activation of Group 1 metabotropic glutamate receptors (mGluRs), FMRP is dephosphorylated and rapidly degraded, allowing for local translation of FMRP-associated mRNAs (Ceman, O’Donnell et al. 2003, Hou, Antion et al. 2006, Nalavadi, Muddashetty et al. 2012).

Dysregulation of mGLuR signaling is thought to play a central role in disease pathogenesis, and both genetic and pharmacologic targeting of these receptors suppresses phenotypes in mice (Bear, Huber et al. 2004, Dolen, Osterweil et al. 2007, Michalon, Sidorov et al. 2012). However, studies of mGluR inhibitors in humans were unsuccessful (Berry-Kravis, Des Portes et al. 2016, Berry-Kravis, Lindemann et al. 2017). Other preclinical studies in *Fmr1* KO mice and drosophila models demonstrated dysfunction in GABAergic signalling (Chang, Bray et al. 2008, Braat, D’Hulst et al. 2015). This too led to a series of clinical trials that failed to meet their primary endpoint (Berry-Kravis, Hagerman et al. 2017, Berry-Kravis, Lindemann et al. 2017, Ligsay, Van Dijck et al. 2017). More recently, FMRP was found to have additional functions in targeting of ion channel proteins in neurons through direct protein-protein interactions, and these functions underlie some of the phenotypic and electrophysiological abnormalities in *Fmr1* KO mice (Brown, Kronengold et al. 2010, Lee, Ge et al. 2011, Deng, Rotman et al. 2013). FMRP also functions as part of the RISC complex in miRNA translational silencing and has poorly understood nuclear functions which may be relevant to disease phenotypes (Cheever and Ceman 2009, Kim, Bellini et al. 2009, Alpatov, Lesch et al. 2014, Korb, Herre et al. 2017). Thus, one potential explanation for the lack of success in human clinical trials to date is that the pleiotropic functions played by FMRP in neurons and other cell types may be difficult to correct with any treatment targeting only one dysregulated pathway(Berry-Kravis, Lindemann et al. 2017).

An alternative approach to therapeutic development in Fragile X Syndrome involves directly targeting the proximal event in disease pathogenesis- the transcriptional silencing of the *FMR1* gene(Tabolacci, Palumbo et al. 2016). While most FXS patients exhibit CGG repeat methylation, in a fraction of cases this methylation is incomplete or absent, allowing for continued *FMR1* transcription (Nolin, Glicksman et al. 1994, Jacquemont, Curie et al. 2011). However, large transcribed repeats still exhibit marked translational inefficiency, presumably due to the repeat element precluding ribosomal scanning through the start codon utilized to generate FMRP (Feng, Zhang et al. 1995). Despite this, in cases where some *FMR1* transcription occurs, expression correlates with both symptom severity and response to therapeutics (Tassone, Hagerman et al. 1999, Jacquemont, Curie et al. 2011). These findings suggest that even small changes in *FMR1* mRNA expression might lead to phenotypic improvements in patients.

Previous work utilizing pharmacological approaches to reactivation of the *FMR1* locus met with some success. Application of non-specific demethylating agents such as 5-azadeoxycytidine (5-azadC) to Fragile X patient derived cells is sufficient to at least transiently enhance *FMR1* transcription and in some cases recover FMRP expression (Chiurazzi, Pomponi et al. 1998). Similarly, treatment with agents that alter the epigenetic landscape, such as the SIRT1 histone deacetylase inhibitor splitomycin, can also re-activate FMR1 transcription in patient derived lymphoblastoid cell lines, suggesting that other epigenetic manipulations may also be effective (Biacsi, Kumari et al. 2008). Approaches coupling these two techniques hold promise at extending the potential effects in patient cells (Kumari and Usdin 2016). However, many of these agents are toxic in humans and have the potential for significant off-target activity elsewhere in the genome, potentially confounding their use clinically in FXS patients.

An important step in developing methods for reactivation of *FMR1* transcription is identifying a model that demonstrates the developmental epigenetic silencing that occurs in FXS patients. Human embryonic stem cells (hESCs) are important disease models for studying developmental processes for which no other suitable models exist. Previous studies in FXS hESC show that some full mutation hESC remain unmethylated following derivation and exhibit gradual loss of *FMR1* mRNA during directed neuronal differentiation (Eiges, Urbach et al. 2007, Telias, Segal et al. 2013, Colak, Zaninovic et al. 2014)), similar to the silencing observed in human FXS fetuses (Malter, Iber et al. 1997). In other lines, however, gene silencing occurs absent differentiation and appears to be repeat-length dependent, with expansions beyond 400 repeats demonstrating greater silencing (Avitzour, Mor-Shaked et al. 2014, Brykczynska, Pecho-Vrieseling et al. 2016, Zhou, Kumari et al. 2016). However, most hESC lines derived and characterized to date are not currently available in the United States for federally funded research.

More recently, researchers have taken a more targeted approach to *FMR1* gene reactivation using the Clustered Regularly Interspaced Palindromic Repeats-CRISPR associated protein 9 (CRISPR-Cas9) system (Doudna and Charpentier 2014). This technique utilizes either one or a set of single guide RNAs (sgRNAs) to target the CRISPR-Cas9 complex to specific genomic loci. The Cas9 endonuclease then nicks the DNA, allowing for either introduction of a deletion or for homology-directed repair. Two separate groups have now utilized CRISPR-Cas9 to delete expanded CGG repeats in Fragile X patient derived cells (Park, Sung et al. 2016, Xie, Gong et al. 2016). In both cases, removal of the repeat led to reactivation of the *FMR1* gene and production of FMRP.

In addition to endonuclease-mediated gene editing, the CRISPR-Cas9 system can also be modified to allow for targeted gene expression modulation in multiple systems (Hsu, Lander et al. 2014). Of particular interest is the use of CRISPR-Cas9 to activate gene expression by using an endonuclease-deficient Cas9 (dCas9) fused to a transcriptional activator (Perez-Pinera, Kocak et al. 2013). Here, we show evidence for the targeted activation of the *FMR1* gene using a dCas9 fused to multiple domains of the VP16 transcriptional activator. Our initial studies in cell lines show differential activity for the various dCas9-VP16 fusion constructs with the most robust activity seen with the dCas9-VP192 construct. This system was used in a newly characterized human embryonic stem cell (hESC) line derived from a Preimplantation Genetic Diagnosis (PGD) FXS embryo. The FXS hESCs show a passage-dependent silencing of the *FMR1* transcript. The dCas9-VP192 construct coupled with guide RNAs targeting the CGG repeat elicited significant activation of *FMR1* transcription in both the early and late passage FXS hESCs and in patient derived Neural Progenitor Cells (NPCs). Overall, these data provide proof-of-principal evidence that CRISPR-dCas9 based transcriptional activation approaches can reactivate *FMR1* transcription even in the setting of large methylated repeats and that targeting the repeat itself may enhance such efforts by providing multiple sequential binding sites for sgRNAs, effectively leveraging the disease mutation to greater efficacy.

## MATERIAL AND METHODS

### CRISPR guide RNA design and plasmids

Promoter-targeted gRNA sequences were identified within 500 nucleotides upstream of the main transcriptional start site based on the prediction of on-target to off-target effect in the human genome and arrangement within the region using the CRISPR design web portal (Hsu, Scott et al. 2013). These sequences and the CGG repeat sequence were cloned into the pSPgRNA plasmid by replacing the sequence between the BbsI sites using the Q5 site-directed mutagenesis kit (NEB) and the primers listed in Suppl.Table 1. All dCas9 expression plasmids were obtained from Addgene. pcDNA-dCas9-VP64, pSPgRNA, and pLV hUbC-dCas9 VP64-T2A-GFP were gifts from Charles Gersbach (Addgene plasmid # 47107, 47108, 53192, respectively)(Perez-Pinera, Kocak et al. 2013). SP-dCas9-VPR was a gift from George Church (Addgene plasmid # 63798) (Chavez, Scheiman et al. 2015). pAC93-pmax-dCas9VP160 was a gift from Rudolf Jaenisch (Addgene plasmid # 48225) (Cheng, Wang et al. 2013). pCXLE-dCas9VP192-T2A-EGFP was a gift from Timo Otonkoski (Addgene plasmid # 69536) (Balboa, Weltner et al.). pcDNA3.1(+) and pEGFP-N1 served as control plasmids.

### Cell culture and transfection of HEK cells

HEK293T cells (ATCC) were maintained at 37°C, 5% CO2 in Dulbecco’s modified Eagle’s Medium supplemented with 10% FBS, 100 U/mL penicillin, and 100 µg/mL streptomycin following standard procedures. Transfections were performed using Lipofectamine 3000 transfection reagent (Invitrogen) according to manufacturer’s instructions. Transfection efficiencies were routinely higher than 80%, as determined by fluorescence microscopy after delivery of a control eGFP expression plasmid. dCas9 expression plasmids were transfected at a mass ratio of 3:1 to either the CGG gRNA expression plasmid or the identical amount of gRNA expression plasmid consisting of a mixture of equal amounts of the four promoter-targeted gRNAs. Cells were harvested 48 h after transfection.

### RNA isolation and quantitative RT-PCR

Total RNA was isolated using the Quick RNA Miniprep kit (Zymo research) with on-column DNase I treatment followed by cDNA synthesis using the iScript Reverse Transcriptase kit (Biorad) according to the manufacturer’s protocol. Quantitative RT-PCR was performed on the Bio-Rad iCycler real-time detection system using the iQ SYBR Green Supermix (Biorad) and the primers (IDT) listed in Supplemental Table 1. Primer specificity was confirmed by gel electrophoresis and melting curve analysis. Relative fold expression for genes of interest were calculated using the comparative CT method (Schmittgen & Livak) with HPRT as the internal control. Technical triplicates were averaged and recorded for each sample. To identify potential off-target genes, a blast search of the human transcriptome was performed with a sequence of 10 CGG repeats (30 nucleotides). The hits were sorted based on their total score. Primers for qPCR were designed for all genes with a score greater than or equal to FMR1. All of these genes contained repeats in the 5′UTR similar to FMR1.

### Western Blot

Cells were washed with cold PBS and lysed on ice in RIPA buffer (50mM Tris-Cl pH-8.0, 150mM NaCl, 0.1% SDS, 0.1% Sodium deoxycholate, 1% NP-40) with complete protease inhibitor cocktail (Roche). Samples were centrifuged at 14,000 RPM for 5 min at 4°C and supernatant was transferred to a clean tube. For western blot, protein lysates were boiled in Laemelli buffer and separated on SDS-PAGE gels. Gels were transferred to PVDF membranes, blocked with 5% nonfat dry milk, and probed with mouse anti-FMRP (6B8) (Biolegend 834601), rabbit anti-Cas9 (Clontech 632607), and rat anti-tubulin (Abcam ab-6160) primary antibodies. Secondary antibodies were goat anti-mouse HRP (Jackson Immunoresearch), IRDye 800 goat anti-rabbit IgG (LI-COR), and IRDye 800 goat anti-rat IgG (LI-COR), respectively. Antibodies were detected using an Odyssey imager or using Western Lightning Plus-ECL substrate (Perkin-Elmer) and developed on film.

### Immunocytochemistry

Cells were cultured as described in chamber slides or on coverslips. The media was removed and cells were washed with 1X PBS and fixed in 4% PFA /4% Sucrose solution for 15 min at room temperature (RT). The cells were permeabilized with 0.1% Triton X-100 for 5 min at RT and were blocked in a 5% Normal Goat Serum solution for 1 hour at RT. The cells were stained with primary antibodies overnight at 4°C followed by three washes in 1X PBS for 5 min each. The antibodies used were: mouse anti-FMRP (6B8) at 1:250 dilution (Biolegend 834601), rabbit anti-Cas9 (Clontech 632607) at 1:150 dilution, anti-MAP2 (Millipore Ab5622) at 1:1000 dilution, SOX2, and PAX6. The cells were then stained with species specific secondary antibodies conjugated to Alexa 488, 568, or 635 fluorophores and mounted using Prolong Gold with DAPI. Images were captured on an inverted Olympus FV1000 laser-scanning confocal microscope.

### ES Cell Line Derivation and Characterization

Human embryos were donated, under two conditions, to MStem Cell Laboratory’s Institutional Review Board (IRB) approved study (HUM00028742) entitled *“Derivation of Human Embryonic Stem Cells”*. First, embryos made for reproductive purposes, not genetically tested, frozen, and no longer required for reproduction were donated (e.g. UM4-6). Second, partners with a known history of familial Fragile-X elected to perform *in vitro* fertilization and preimplantation genetic diagnosis (PGD), irrespective of embryo donation, to reduce the risk of having a child with Fragile X-spectrum disorder. The female partner was an FMR1 pre-mutation carrier with a mutant allele determined to have 108 and 115 CGG repeats on two separate evaluations. The female partner had three paternal uncles with Fragile X-spectrum disorder. *In vitro* produced embryos were biopsied as blastocysts on day 5 of development, and trophectoderm cells were genetically assessed by an off-site genetic analysis company. Blastocysts were vitrified and cryo-stored until PGD results were obtained. Embryos with PGD results showing the mutant maternal haplotype and no paternal X chromosome (affected male) where consented for donation and shipped to MStem Cell Laboratory (e.g. UM139-2).

Following hESC production and characterization, documents demonstrating adherence to NIH-established guidelines for embryo donation and hESC production of UM4-6 and UM139-2 were submitted to NIH for placement on the NIH hESC Registry and approvals were granted on 02/02/2012 (Registration # -0147) and 09/29/2014 (Registration # -0292), respectively. Derivation of hESCs and their derivatives prior to acceptance on the NIH registry were performed with non-federal funds. Additionally, studies after placement on the NIH registry were also supported by non-federal funds. Briefly, blastocyst morphology was assessed four hours after embryos were warmed, and dictated the mode of hESC derivation. Laser-dissected inner cell masses were plated on human foreskin fibroblast (HFF)-feeders to obtain early hESC colonies that were manually split after 5 to 7 days and expanded on HFF for establishment and characterization of hESC lines. HESC lines were tested for pluripotency marker expression (Oct4, Nanog, Sox2) by Q-PCR and protein expression by immunofluorescence (Oct4, Nanog, Sox2, SSEA4 and TRA-1-60). hESCs were differentiated for 21 days in culture as embryoid bodies and tested for expression of lineage markers by Q-PCR of endoderm (α-fetoprotein and GATA4), mesoderm (brachyury and VE-Cadherin) and ectoderm (TUJ-1 and Krt-18). Finally, G-band karyotyping of UM4-6 and UM139-2 demonstrated euploid hESC lines.

### Culture and transfection of hESCs

Undifferentiated hESCs were cultured in mTeSR1 media (Stem Cell Technology) on MatriGel-coated plates with daily media changes and were passaged at 1:5 to 1:10 using L7 passaging media (Lonza) or 1 mM EDTA. For transfections, undifferentiated hESCs were plated as small colonies on MatriGel-coated plates in mTeSR1 media containing 10 µM Rock Inhibitor and grown overnight. Media was replaced with mTeSR1 the next day. Cells were allowed to recover for at least 4 hours and media was replaced again just prior to transfections. Transfections were performed using plasmids as described above and TransIT LT1 transfection reagent (Mirus Bio) according to manufacturer’s instructions. Cells were cultured with daily media changes and harvested 48 h after transfection for RNA isolation and 72 h after transfection for western blots.

### Southern Blot

Southern blotting was performed as in (Gold, Radu et al. 2000) with modifications. Briefly, genomic DNA was isolated using the Qiagen DNeasy Blood and Tissue kit. 10ug of genomic DNA from each cell line was digested with HindIII and EagI overnight. The digoxigenine (DIG)-labeled probe was amplified from the pE5.1 plasmid, using forward primer: CGCCAAGAGGGCTTCAGGTCTCCT and reverse primer: GAGACTGTTAAGAACCTAAACGCGGG. The digested genomic DNA was resolved on a 0.7% agarose gel prior to southern blotting. The nylon membrane was processed using the commercially available DIG Easy Hyb solution and DIG Wash and Block Buffer Set (Roche). DIG was antibody labeled with Anti-digoxigenine-AP, Fab fragments (Sigma), processed using CDP-Star substrate (ThermoFisher), and detected on film. A wild-type band (~30 repeats) in Control lines appears at ~2.8kb, whereas the expanded and methylated repeat in the Fragile X line appears at ~7.6kb (800 repeats) which is ~2.4kb above where a non-expanded, methylated DNA fragment would appear (~5.2kb).

### Methylation qPCR

Genomic DNA was isolated from cell lines using the Qiagen DNeasy Blood and Tissue Kit. 2ug of DNA from each was bisulfite converted using the EpiTect Bisulfite kit (Qiagen). qPCR was performed on 40ng of bisulfite converted DNA using the iQ SYBR Green supermix (BioRad). Three primers were used per sample, in triplicate. *FMR1* methylation-specific primers, forward: GGTCGAAAGATAGACGCGC and reverse: AAACAATGCGACCTATCACCG; *FMR1* unmethylated-specific primers, forward: TGTTGGTTTGTTGTTTGTTTAGA and reverse: AACATAATTTCAATATTTACACCC; and primers for the housekeeping gene CLK, which should not undergo CpG methylation in the region of amplification, forward: CGGTTGATTTTGGGTGAAGT and reverse: TCCCGACTAAAATCCCACAA. Amplification of both methylated and unmethylated *FMR1* was normalized to the housekeeping gene, then a ratio was created using the two values. In the Control fibroblasts where no amplification was detected with the methylation-specific primers, methylation was set at 0%.

### Directed differentiation of hESC to NPCs and Neurons

Neural induction was performed using a dual-SMAD inhibition(Shi, Kirwan et al. 2012) protocol with modifications. In brief, undifferentiated hESCs in two wells of a 6-well plate were grown to approximately 80% confluence, dissociated with EDTA, and plated into a single well of a MatriGel-coated 6-well plate with TeSR-E8 containing 10 µM Rock Inhibitor (Y-27632). The cells were confluent the next day and neural differentiation was induced using neural maintenance media (referred to here as 3N) containing 1 µM dorsomorphin and 10 µM SB431542. The cells were cultured for 12-14 days with daily media changes. Neuroepithelial sheets were then combed into large clumps, passaged, and maintained on MatriGel-coated plates in rosette media (3N containing 20 ng/ml FGF2) with daily media changes until neural rosettes appeared. Rosettes were manually picked and dissociated into single cells using Accutase. NPCs were plated onto MatriGel-coated plates, grown in neural expansion media (3N containing 20 ng/ml FGF and 20 ng/ml EGF) with media changes every other day, and passaged as needed using Accutase. For differentiation into neurons, NPCs were plated at a density of approximately 1.5 x 10^5^ cells/mL in neural expansion media on PLO-laminin coated plates or coverslips, allowed to grow for 24 hours, and switched to neural maintenance media. Neurons were maintained for up to 6 weeks with half media changes every other day and a full media change supplemented with 1 µg/ml laminin every 10 days.

### RNA Sequencing and GO Analysis

Sequencing was performed by the UM DNA Sequencing Core, using the Illumina Hi-Seq 4000 platform, single end, 50 cycles, mRNA prep. At the UM Bioinformatics Core, files from the Sequencing Core’s storage were concatenated into a single fastq file for each sample. The quality of the raw reads data for each sample was checked using FastQC (http://www.bioinformatics.bbsrc.ac.uk/projects/fastqc/) (version 0.11.3) to identify features of the data that may indicate quality problems (e.g. low quality scores, over-represented sequences, inappropriate GC content). The Tuxedo Suite software package was used for alignment, differential expression analysis, and post-analysis diagnostics (Langmead, Trapnell et al. 2009, Trapnell, Pachter et al. 2009, Trapnell, Hendrickson et al. 2012). Briefly, the reads were aligned to the reference mRNA transcriptome (hg 19)(http://genome.ucsc.edu/) using TopHat (version 2.0.13) and Bowtie2 (version 2.2.1.). Default parameter settings for alignment were used, with the exception of: “--b2-very-sensitive” telling the software to spend extra time searching for valid alignments. FastQC was used for a second round of quality control (post-alignment), to ensure that only high quality data would be input to expression quantitation and differential expression analysis. Cufflinks/CuffDiff (version 2.2.1) was used for expression quantitation, normalization, and differential expression analysis, using hg19.fa as the reference genome sequence. For this analysis, the parameter settings were: “--multi-read-correct” to adjust expression calculations for reads that map in more than one locus, as well as “--compatible-hits-norm” and “--upper-quartile–norm” for normalization of expression values. Diagnostic plots were generated using the CummeRbund R package. Locally developed scripts were used to format and annotate the differential expression data output from CuffDiff. Briefly, genes and transcripts were identified as being differentially expressed based on three criteria: test status = “OK”, FDR ≤ 0.05, and fold change ≥ ± 1.5. Genes and isoforms were annotated with NCBI Entrez GeneIDs and text descriptions. iPathwayGuide (Advaita Corporation-http://www.advaitabio.com) was used to model the biological relevance of differentially expressed genes for each algorithm, as well as a meta-analysis comparing the two approaches.

## RESULTS

### Transcriptional activation of the *FMR1* gene by CRISPR-dCas9 fused to VP16 activation domains

To determine whether use of CRISPR targeted transcriptional activators could augment FMR1 expression, we first tested them in HEK 293T cells that have a normal sized (23) CGG repeat in the 5’UTR of the *FMR1* gene. We designed multiple guide RNAs (gRNAs) to the promoter region or to the CGG repeat of the *FMR1* gene. These gRNAs were used along with the catalytically-inactive dCas9 fused to different versions of the VP16 transcriptional activation domain (Fig. 1A). At 48 hours post transfection, we observed a significant increase in *FMR1* transcript levels using the dCas9-VP64 construct with both the promoter pool and CGG gRNAs (Fig. 1B) compared to a scrambled control gRNA or to cells transfected with only GFP. We next compared activation efficiencies for both sets of gRNAs with different versions of dCas9 fused with either the chimeric activation domain VPR (composed of the activation domains of VP64, p65 and Rta linked in tandem), or multiple domains of VP16 (Fig. 1C). The strongest transcriptional activation in heterologous cells was achieved with a CGG repeat-targeted dCas9-VP192, which yielded approximately an 8-fold increase in *FMR1* mRNA levels (Fig. 1C, 1D). The CGG repeat targeted guide robustly increased *FMR1* transcript levels compared to the promoter-pool gRNAs, suggesting that its repetitive binding sites augment the targeting strategy (Fig. 1D).

**Figure 1:**
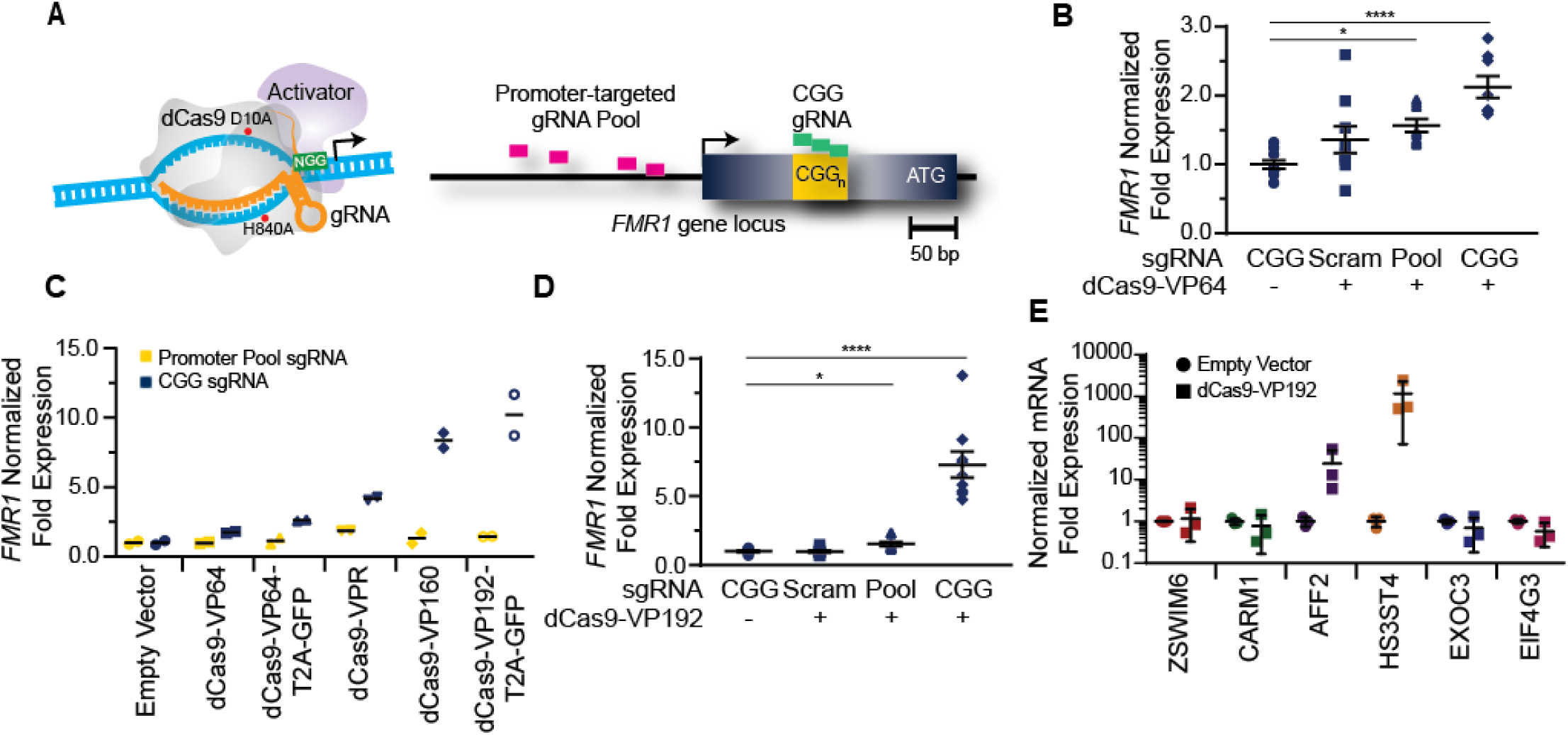
*FMR1* mRNA increases with CRISPR-mediated targeting of transcriptional activators to either the *FMR1* promoter or the CGG repeat. A) Illustration of nuclease-inactive Cas9 (dCas9) fused to a transcriptional activator (left) and guide RNA (gRNAs) targeting regions within the *FMR1* promoter or the CGG repeat (right). The promoter-targeted gRNA pool (pink) consisted of four gRNAs with unique targeting sequences, while the CGG gRNA (green) represents a single targeting sequence capable of tiling across the CGG repeat. B) Relative *FMR1* mRNA expression in HEK293T cells transfected with empty vector or dCas9 fused to VP64 (dCas9-VP64) and non-targeting guide RNA (Scram), a pool of four guide RNAs within the *FMR1* promoter (Pool), or a single CGG repeat guide (CGG). C) Relative *FMR1* mRNA expression in HEK293T cells transfected with an empty vector or dCas9 “second generation” activators and the *FMR1* promoter gRNAs or the CGG gRNA. D) Relative *FMR1* mRNA expression in HEK293T cells transfected with a control plasmid or dCas9 fused to VP192 (dCas9-VP192) and the indicated gRNA. (For panels B-D, *p<0.05, ****p< 0.0001, Kruskal-Wallis one-way ANOVA with Dunn’s multiple comparison test). E) Relative expression of select CGG repeat-containing genes after transfection of HEK293T cells with CGG gRNA and dCas9-VP192 constructs. (***p<0.001, two-way ANOVA with Sidak’s multiple comparisons test.) For all scatter plots shown, each data point represents an individual well.

Because CGG tandem microsatellites are not unique within the genome, we also queried 6 candidate genes with CGG repeats in their 5’UTR for off target effects in HEK293T cells. We observed an increase in transcript levels for the *AFF2* gene (also called *FXR2*) and *HS3ST4* gene suggesting potential off-target effects in this cell type with a repeat-targeting strategy (Fig. 1E). However, the effects on AFF2 are potentially interesting clinically. Expansion of this CCG repeat triggers hypermethylation and transcriptional silencing of the AFF2 gene, in a fashion that is quite similar to FXS (also known as FRAXA). This results in FRAXE, a rare genetic form of autism and intellectual disability(Knight, Flannery et al., Gecz, Gedeon et al. 1996). Together, these data demonstrates that the dCas9-VP192 system can effectively activate transcription of the *FMR1* gene. Additionally, CGG gRNA provide more robust activation compared to promoter pool gRNAs but with a greater potential for off-target effects.

### dCas9-VP192 increases FMRP levels in HEK293T cells

We next determined if the observed transcriptional changes correlated with enhanced production of FMRP. FMRP levels were measured in HEK293T cells transfected with either the promoter pool or CGG-repeat targeted gRNAs and the dCas9-VP192 construct. By immunocytochemistry, cells transfected with CRISPR constructs show an increase in FMRP signal with either CGG or promoter targeted gRNAs, but not with scramble guide RNA(Fig. 2A). Western blot analysis of transfected cells demonstrated a significant increase in FMRP protein in CGG repeat targeted gRNAs compared to control transfections (Fig. 2B and 2C). Thus, targeted activation of the *FMR1* gene using a dCas9-VP192 system increases both *FMR1* mRNA and FMRP levels in human cells at normal repeat sizes.

**Figure 2:**
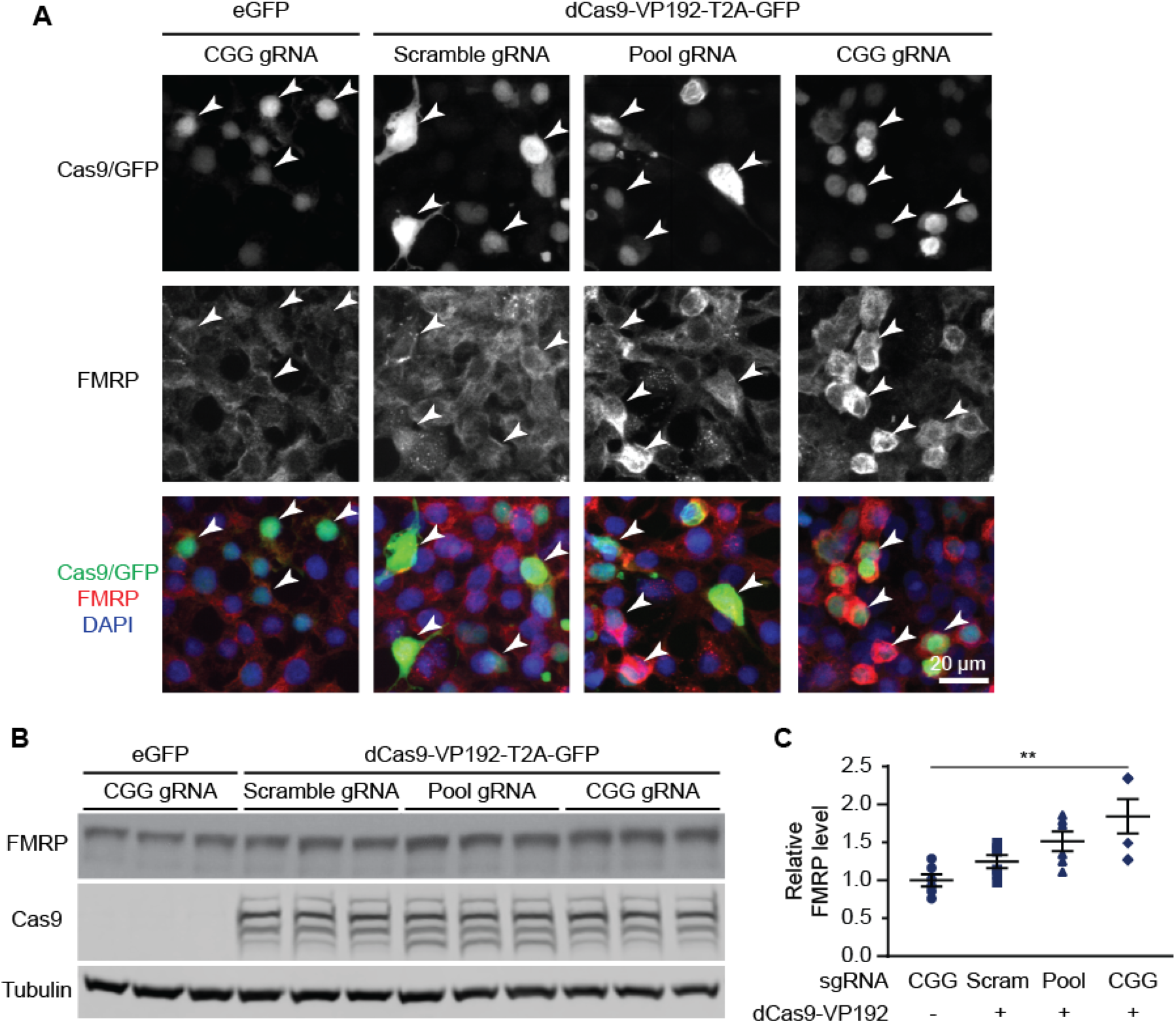
CRISPR-mediated transcriptional activation increases FMRP protein abundance. A) Immunocytochemistry of HEK293T cells transfected with eGFP or dCas9-VP192 and gRNAs, as indicated. Single channel and merged images are shown with Cas9/GFP (green), FMRP (red), and DAPI (blue). White arrowheads indicate transfected cells. Scale bar represents 20 μm. B) Western blots showing triplicate samples of HEK 293T cells transfected with control plasmid (eGFP) or with dCas9-VP192 and indicated gRNAs and immunoblotted for FMRP, Cas9 and Tubulin. C) Quantification of western blots from HEK293T cells transfected as indicated. Data are shown as FMRP normalized to tubulin and relative to the control plasmid. (**p< 0.01, Kruskal-Wallis one-way ANOVA with Dunn’s multiple comparison test.)

### FXS hESCs exhibit passage-dependent silencing of *FMR1* prior to neuronal differentiation

An important first step in developing a method for reactivation of *FMR1* transcription is identifying a model that recapitulates the developmental epigenetic silencing that occurs in FXS patients. Until recently, the UM139-2 PGD hESC line was the only Fragile X hESC line on the NIH registry of approved lines for federally-funded research in the United States (https://stemcells.nih.gov/research/registry.htm). However, different Fragile X hESC lines exhibit variability in terms of their methylation and *FMR1* transcription (Avitzour, Mor-Shaked et al. 2014). We therefore characterized this new FXS hESC line.

The embryo from which this hESC line (UM 139-2) was derived was determined to be affected with Fragile X Syndrome through preimplantation genetic diagnosis (PGD). This blastocyst was cryopreserved after testing and sent to MStem Cell laboratories, where derivation of hESCs took place (Fig. 3A-C). Pluripotency and fidelity of this line was confirmed by RT-PCR and immunohistochemistry for pluripotency markers (Fig. 3D, E). The line was capable of embryoid body formation containing all 3 germ layers, consistent with pluripotency (Fig. 3F). DNA fingerprinting and karyotyping demonstrated a 46 XY euploid genetic background (Fig. 3G).

**Figure 3:**
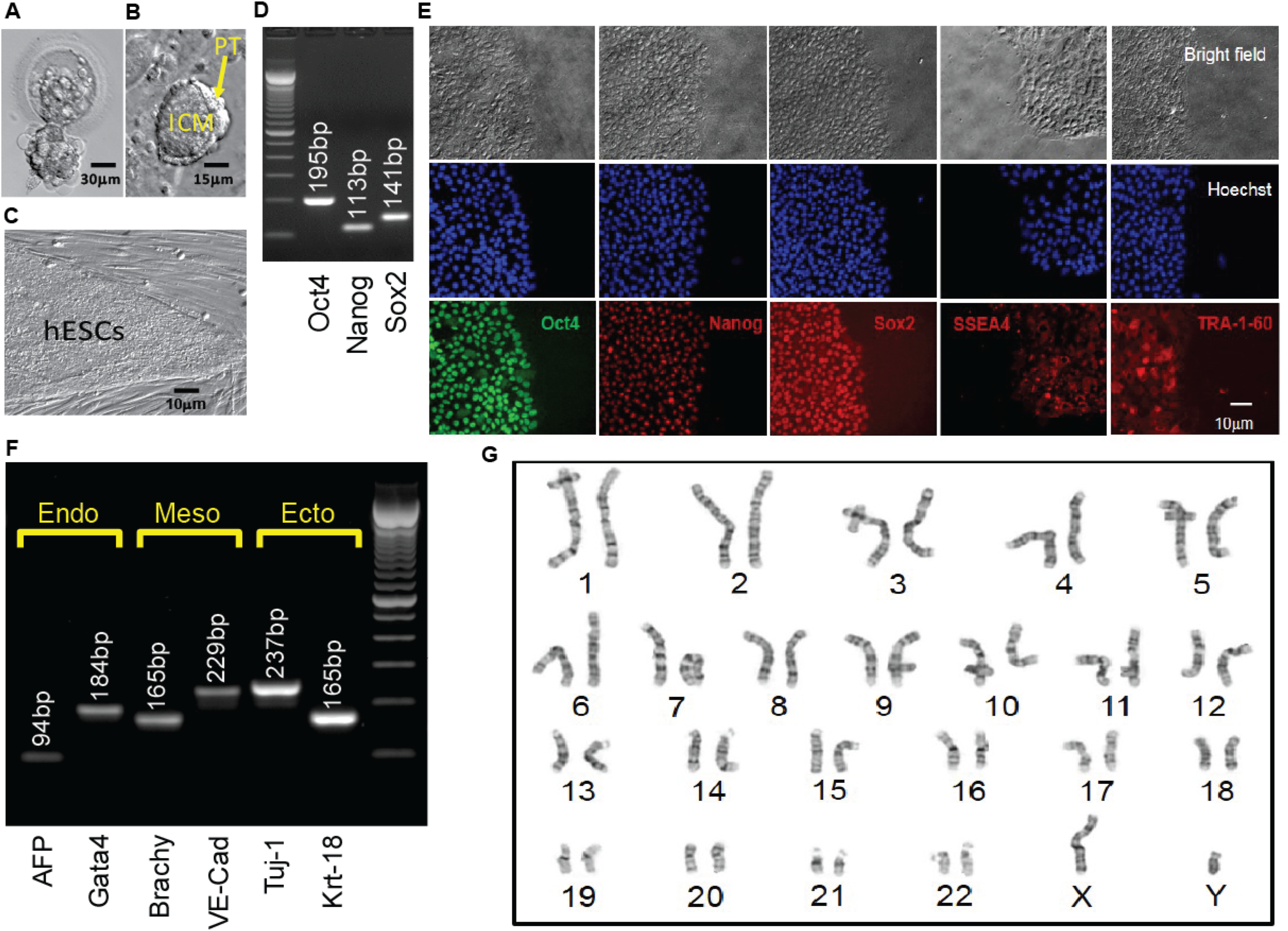
Derivation, expansion, and characterization of Fragile X-disease specific human embryonic stem cell (Frag-X-ds-hESC; UM139-2 PGD) line. (A) Human blastocyst with *FMR1* expansion that was cryopreserved, donated, shipped, and warmed prior to attempting hESC derivation. Scale bar represents 30 μm. B) The inner cell mass (ICM) with surrounding polar trophectoderm (PT) was laser-dissected from the blastocyst and plated/attached on human foreskin fibroblast (HFF). This micrograph represents the early Frag-X-ds-hESC colony before the first passage, 5 days after laser-dissection and plating of the ICM/PT (P0D5) Scale bar represents 15μm. C) Expanding undifferentiated Frag-X-ds-hESCs with tight colony borders on a human foreskin fibroblast feeder layer (P3D3). Scale bar represents 10μm. D) Undifferentiated Frag-X-ds-hESCs expressed pluripotency markers (Oct4, Nanog, Sox2) as assessed by qPCR. Electrophoresis demonstrated anticipated amplicons for each pluripotency marker PCR primer sets. E) Expanded undifferentiated Frag-X-ds-hESCs with tight colony borders on Martigel (brightfield micrographs) expressed pluripotency marker proteins in the nucleus (same location as Hoechst staining; Oct4, Nanog, Sox2) or cytoplasmic/cell membrane associated (SSEA4 and TRA-1-60). F) Frag-X-ds-hESCs were differentiated into embryoid bodies for 21 days in culture. Differentiated Frag-X-ds-embryoid bodies expressed linage marker RNA of endoderm [α-fetoprotein (AFP) and GATA4], mesoderm [brachyury (Brachy) and VE-Cadherin (VE-Cad)] and ectoderm [neuron-specific class III beat-tubulin (Tuj-1) and keratin-18 (Krt-18)] with anticipated amplicon size by electrophoresis for each linage marker PCR primer set. Scale bar represents 10 μm (G) at passage 6 undifferentiated Frag-X-ds-hESCs were sent to Cell Line Genetics (Madison, WI) for G-Band karyotyping and reported to be a 46XY, euploid hESC line.

We next characterized the line in terms of its Fragile X mutation. Southern blot analysis indicated that this hESC line contains a Fragile X full mutation with approximately 800 CGG repeats (Fig. 4A). The first characterized FXS hESC line, HE-FX, exhibited no methylation in the hESC state, but instead demonstrated methylation and transcriptional silencing with cellular differentiation (Eiges, Urbach et al. 2007). However, more recent studies suggest that this property is not universal, with some Fragile X hESCs exhibiting early methylation and silencing. To evaluate whether the repeat was methylated in in UM 139-2 hESC line, we performed methylation-specific quantitative-PCR on early passage (<20 passages) FXS hESCs. This demonstrated significant, although incomplete, methylation over the *FMR1* promoter region compared to control hESCs and FXS fibroblasts (Fig. 4B).

**Figure 4:**
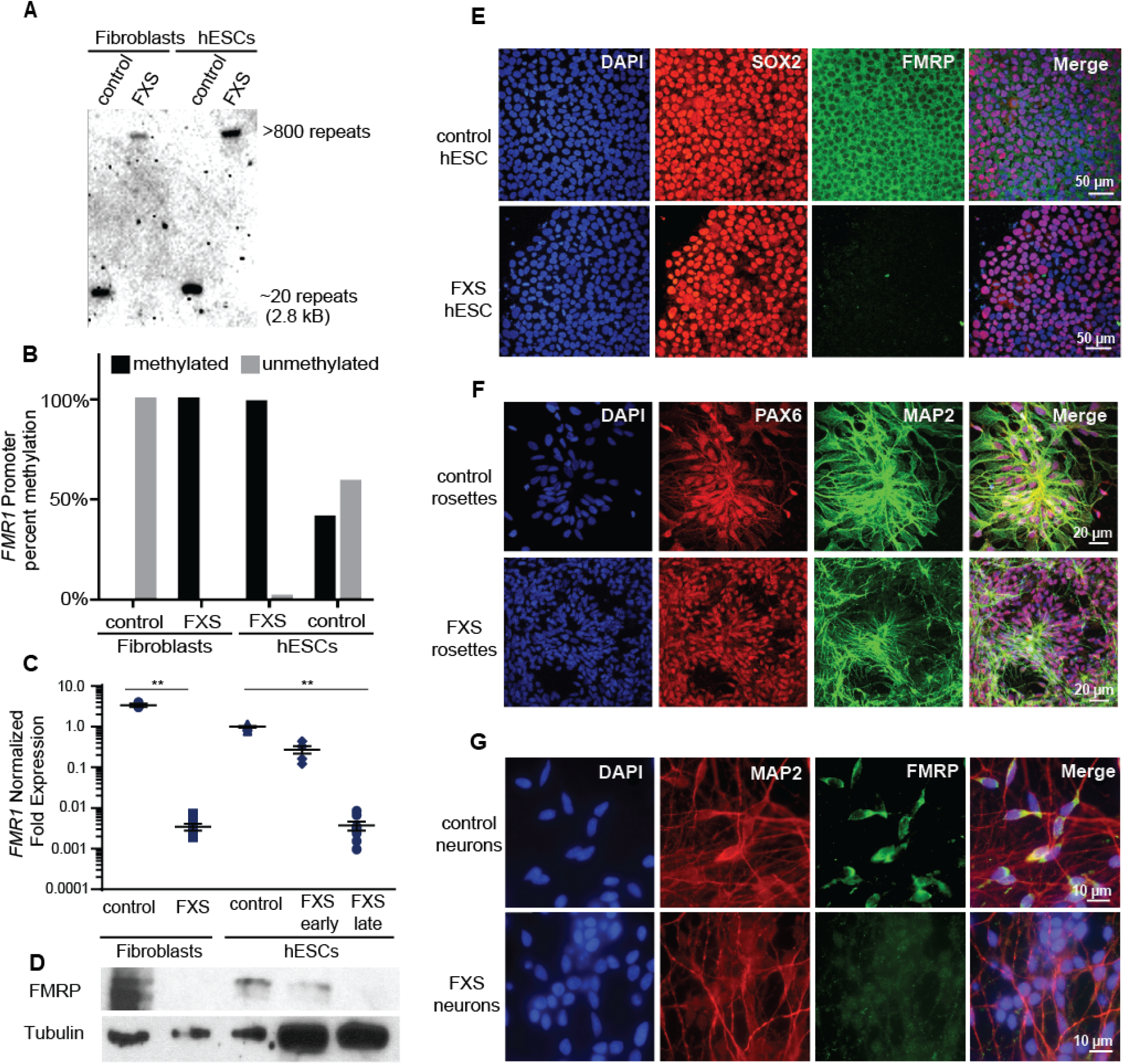
Fragile X hESC line (UM139-2 PGD) carries a large CGG repeat and undergoes spontaneous transcriptional silencing. A) Southern blot using an *FMR1*-specific DIG-labeled probe shows CGG repeat length and methylation status for genomic DNA from control and FXS patient-derived fibroblasts and hESCs. Repeat size is estimated at <800. B) Bisulfite-qPCR using methylation-specific primers reveals the percent methylation at the *FMR1* promoter for genomic DNA from indicated cells. C) Relative *FMR1* transcript levels in control and FXS patient-derived fibroblasts and hESCs. FXS hESCs were assessed at early passages (P13-20) and late passages (P30+). Each data point represents a separate plate of cells and mean with error bars (SEM) is shown for each condition. (**p<0.01, Kruskal-Wallis one-way ANOVA with Dunn’s multiple comparison test). D) Western blots showing FMRP and tubulin in control and FXS patient-derived fibroblasts as well as control hESC and early and late passage FXS hESCs. One-tenth of the lysate was loaded for control fibroblasts and control hESCs. E) Undifferentiated control and FXS hESC colonies immunostained for FMRP (green), pluripotency marker SOX2 (red), and DAPI (blue). Scale bars represent 50 µm. F) Neural rosettes derived from control and FXS hESCs with neuronal lineage marker MAP2 (green), neuroectoderm maker PAX6 (red), and DAPI (blue) immunostaining. Scale bars represent 20 µm. G) Neurons derived from control and FXS hESCs shown with FMRP (green), MAP2 (red), and DAPI (blue) immunostaining. Scale bars represent 10 µm.

To determine the impact of this methylation on FMR1 transcriptional activity, we measured *FMR1* mRNA expression by qRT-PCR. This demonstrated a passage-dependent shift in expression in FXS hESCs. At early passages, *FMR1* mRNA levels were only modestly decreased (30%) compared to controls. However, after continued passages (typically>30 passages, with some variability), *FMR1* mRNA levels became nearly undetectable (0.4% of control levels, Fig. 4C). We observed a similar passage-dependent change in FMRP protein level, although there was a significant deficit in FMRP expression even at early passage numbers, perhaps due to translational blockade (Fig. 4D) (Feng, Zhang et al. 1995, Iliff, Renoux et al. 2013).

To confirm that the absence of FMRP does not preclude differentiation into neurons from FXS hESCs, we performed directed neuronal differentiation using a dual smad inhibitor-based differentiation protocol (Shi, Kirwan et al. 2012). This method successfully produced hESC-derived PAX-6 and MAP2 positive neural rosettes and FXS neurons (Fig. 4 E, F and G). As reported previously for other FXS lines, we also observed a slight delay in neural rosette formation as well as a lower density of neurons from the UM 139-2 FXS line (Telias, Segal et al. 2013). Combined, these data suggest that UM 139-2 FXS hESCs are a good model for investigating methods of reactivating *FMR1* transcription, and the feature of time-dependent transcriptional silencing allows for targeting of reactivation at expanded repeats in different epigenetic contexts.

### dCas9-VP192 activates FMR1 transcription in FXS hESCs and NPCs

Based on our success using dCas9-VP192 to activate transcription of the *FMR1* gene in HEK293T cells, we first tested the same constructs and gRNAs in control hESCs. Control hESCs showed a significant increase in *FMR1* transcript levels and protein levels using the promoter targeted gRNAs with dCas9-VP192 only (Fig. 5A), although the effects were more variable and less robust than those observed in HEK293T cells. We next evaluated whether this increase in mRNA was associated with changes in protein expression. By immunohistochemistry, there was a clear relationship between cells expressing the dCAS9 construct and an increase in FMRP expression for the promoter pool gRNAs but not the CGG repeat gRNAs (Fig 5 B). By western blot as well, only the promoter pool targeted gRNAs demonstrated a significant change in protein expression (Fig. 5 C, D). This discrepancy may reflect differences in efficiency of translation and expression from these vectors between HEK293T cells and hESCs.

**Figure 5:**
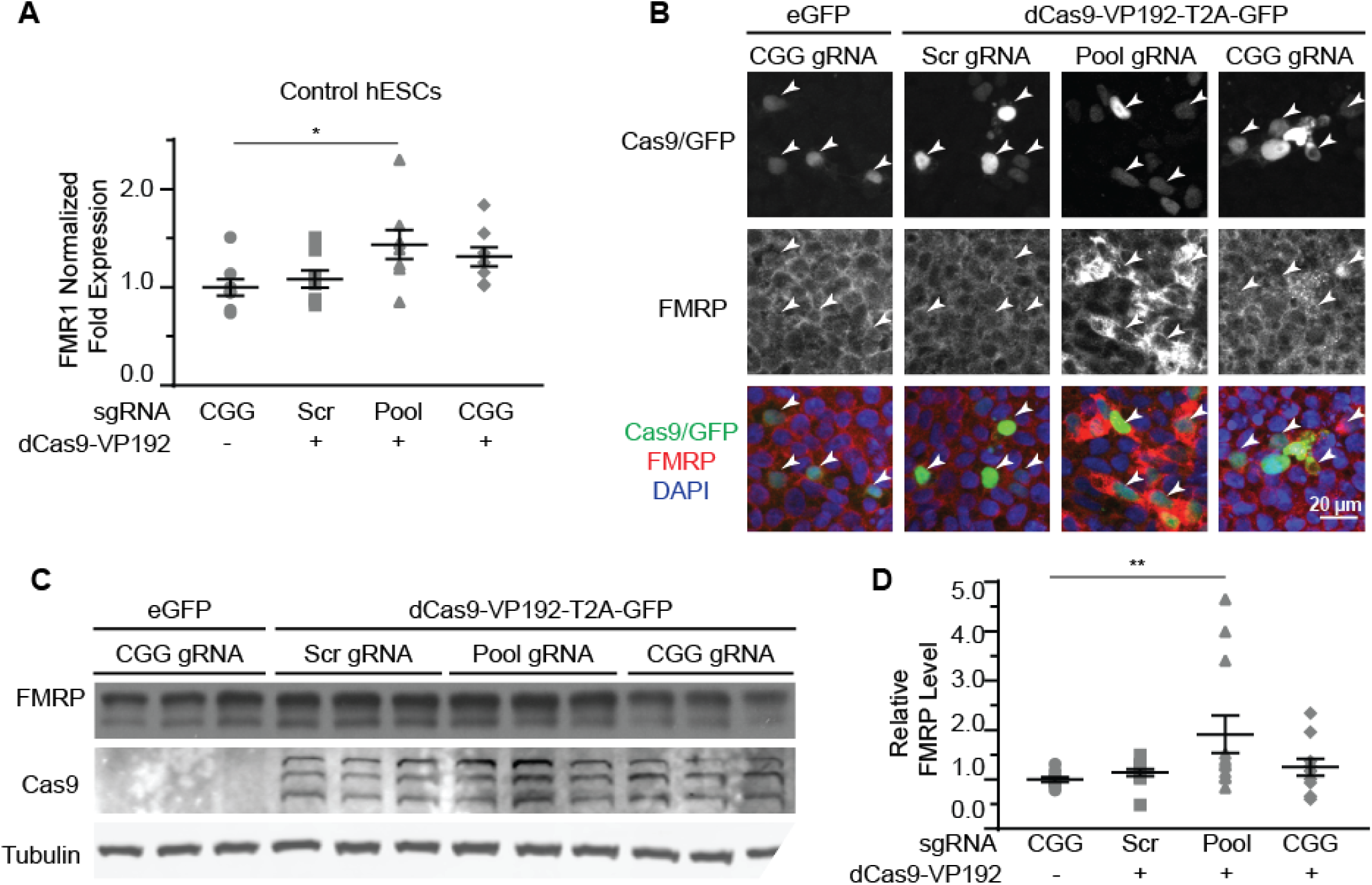
CRISPR-mediated activation enhances *FMR1* transcription and FMRP protein abundance in control hESCs. A) Relative *FMR1* mRNA expression in control (UM4-6) hESCs transfected with a control plasmid or dCas9-VP192 and the indicated gRNAs. (*p<0.05, Kruskal-Wallis one-way ANOVA with Dunn’s multiple comparison test.) B) Images of control hESCs transfected with eGFP or dCas9-VP192 and gRNAs, as indicated. Single channel and merged images are shown with Cas9/GFP (green), FMRP (red), and DAPI (blue). Arrowheads indicate transfected cells. Scale bar represent 20 μm. C) Western blots showing triplicate samples of control hESCs transfected and immunoblotted as indicated. D) Quantification of western blots from control hESCs transfected as indicated. Data are shown as FMRP levels normalized to tubulin and relative to the control plasmid. (** indicates p = 0.0061 by Kruskal-Wallis one-way ANOVA with Dunn’s multiple comparison test.) For all scatter plots, each data point represents an individual well. Data were obtained from three (A) or four (D) independent experiments. The mean with error bars (SEM) is shown for each condition.

We next tested whether the dCas9-VP192 system could re-activate or enhance transcription from the *FMR1* locus in UM139-2 FXS hESCs. Because of their baseline differences in *FMR1* transcription, we evaluated both early and late passage hESCs. In early passage FXS hESCs, both the promoter and CGG gRNAs elicited a 1.3 fold and a 1.8 fold increase, respectively in *FMR1* transcript levels compared to the scrambled guide RNA in the same line (Fig. 6A). However, this increase was significantly greater with the CGG guide RNAs compared to the promoter targeting gRNAs (Fig 6A).

**Figure 6:**
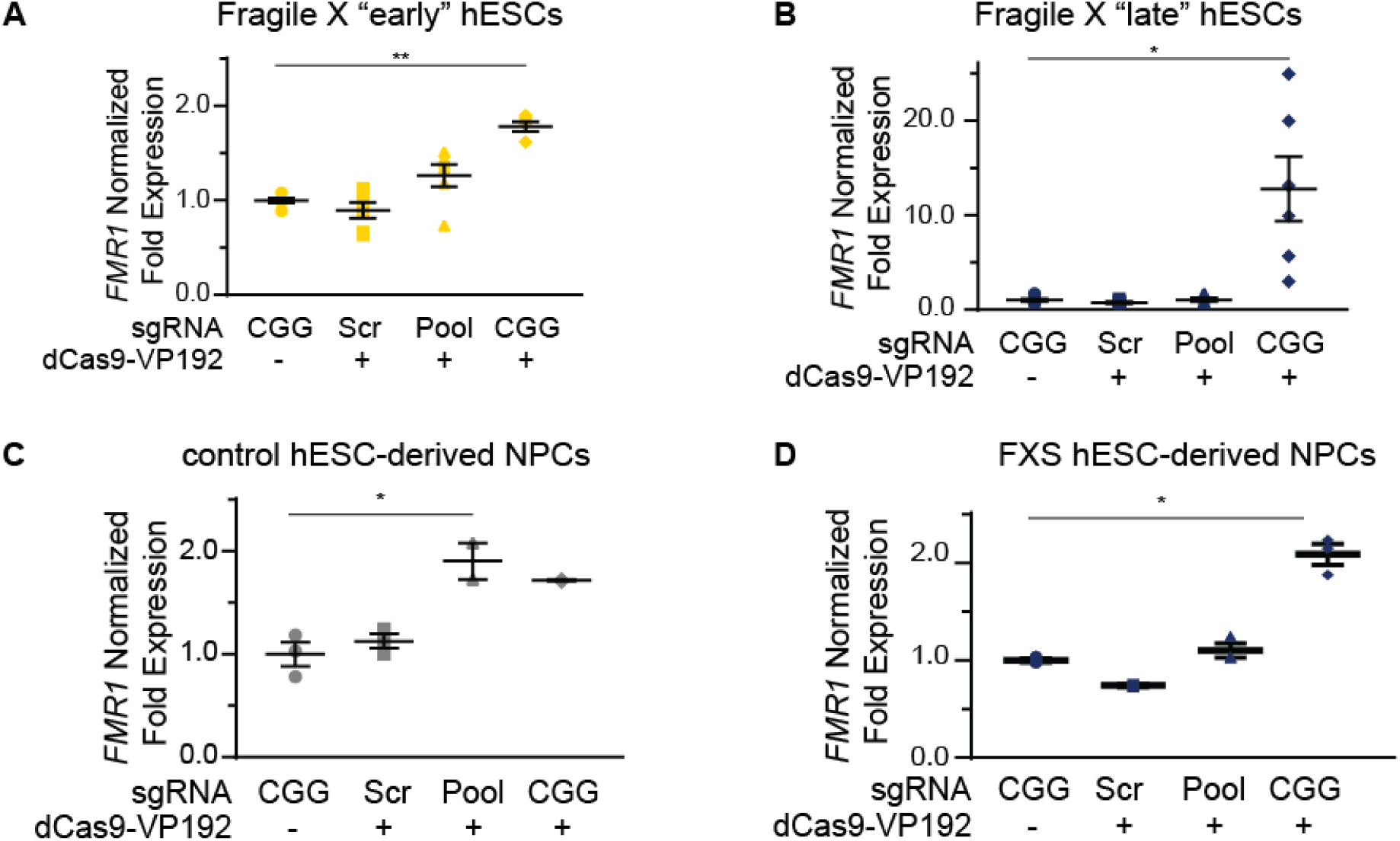
Targeting CRISPR-dCAS9-VP192 to the CGG repeat overcomes transcriptional silencing and selectively enhances transcription of *FMR1* in FXS hESCs and NPCs. A) Relative *FMR1* mRNA expression in early (P23-25) passage FXS hESCs transfected with control plasmid or dCas9-VP192 and the indicated gRNA. (**p<0.01, *p<0.05, Kruskal-Wallis one-way ANOVA with Dunn’s multiple comparison test.). B) Relative *FMR1* mRNA expression in late (P53-57) passage FXS hESCs transfected with control plasmid or dCas9-VP192 and the indicated gRNA. (**p<0.01, *p<0.05, Kruskal-Wallis one-way ANOVA with Dunn’s multiple comparison test.). C) Relative *FMR1* mRNA expression in control hESC-derived NPCs transfected with control plasmid or dCas9-VP192 with the indicated gRNAs. (*p<0.05, Kruskal-Wallis one-way ANOVA with Dunn’s multiple comparison test). D) Relative *FMR1* mRNA expression in late FXS hESC-derived NPCs transfected with control plasmid or dCas9-VP192 with the indicated gRNAs. (*p<0.05, Kruskal-Wallis one-way ANOVA with Dunn’s multiple comparison test).

In late passage FXS hESCs *FMR1* mRNA levels were very low basally-effectively at the level of detection (Fig 4C). Treatment with scrambled gRNA or promoter targeted gRNAs in the setting of dCas9-VP192 had no impact on *FMR1* RNA expression. However, CGG targeted gRNA coupled with dCas9-VP192 led to a marked increase in *FMR1* mRNA expression-upwards of 20 fold in some samples (Fig. 6B). Thus, at both a partially and completely transcriptionally silenced CGG full mutation locus, we observed that targeting a transcriptional activator directly to the repeats elicited the greatest enhancement of *FMR1* mRNA expression. Next, we differentiated the control and late FXS hESCs to NPCs and tested for *FMR1* transcript levels after treating them with the dCas9-VP192 and gRNAs. The NPCs differentiated from the late FXS hESCs were selected since they did not have any baseline *FMR1* transcription, which reflected the disease state more closely. Similar to our observations in undifferentiated hESCs, control NPCs showed a statistically significant increase in *FMR1* levels using the promoter pool targeted and CGG targeted gRNAs (Fig. 6C) while the FXS NPCs showed the same increase in transcript level with the CGG gRNA only (Fig. 6D). Thus, the effects of specific gRNAs with dCas9-VP192 on FMR1 mRNA expression are different in the setting of large CGG repeat expansions across cell differentiation states.

### CGG repeat targeted gRNA shows minimal off target effects in FXS hESCs

We next evaluated whether there were off-target effects elicited by the CGG repeat targeted gRNAs. We first queried the 6 candidate genes identified in HEK293T cells (Fig. 1E). Unlike the case in HEK293T cells, we saw no increase in their mRNA levels in FXS hESCs expressing CGG gRNA and dCas9-VP192 (Fig. 7A). To evaluate for other off-target effects elicited by expression of CGG gRNA and dCas9-VP192, we performed RNA-seq analysis of FXS hESCs treated with scramble gRNA versus CGG gRNA and dCas9-VP192 (Fig. 7B and C). A comparison between FXS hESCs treated with scrambled or CGG gRNA and dCas9-VP192 showed only 35 genes out of 23394 that were differentially expressed between these two conditions (Fig. 7C).

**Figure 7:**
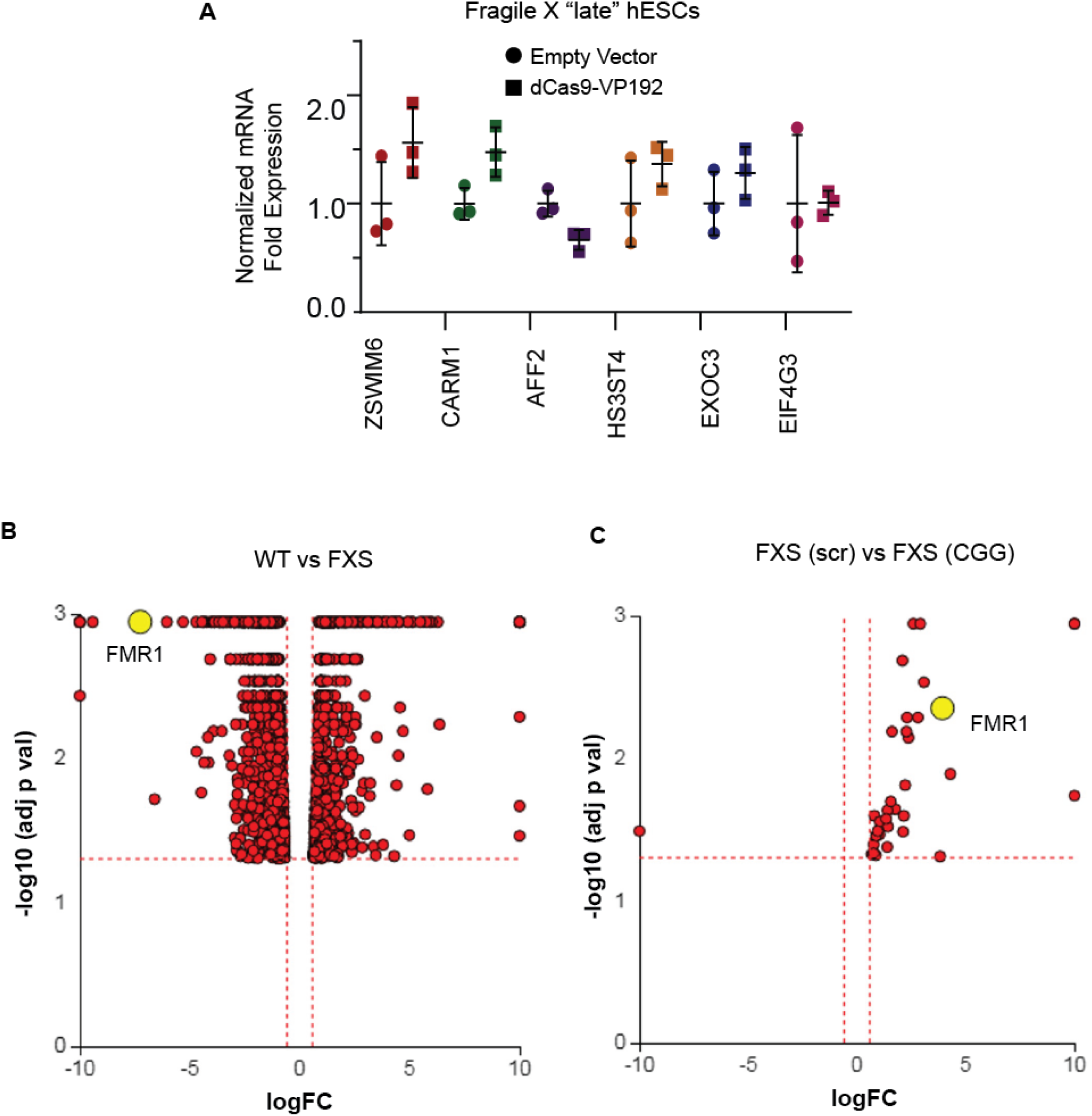
Targeted activation of *FMR1* by CGG guide RNAs has minimal off target effects compared to scramble guide RNA. A) Relative fold expression of select CGG repeat-containing genes after transfection of late passage FXS hESCs with CGG gRNA and eGFP (empty vector) or dCas9-VP192 constructs. B) Volcano plot showing RNA-seq analysis of WT and FXS hESCs. All 1784 significantly differentially expressed (DE) genes are represented in terms of their measured fold change (x-axis) and the significance represented as the negative log (base 10) of the p-value (y-axis). The yellow dot shows the position of *FMR1* mRNA. C) Volcano plot showing RNA-seq analysis of FXS (scramble gRNA treated) and FXS (CGG gRNA treated) late passage hESCs expressing dCas9VP192. All 35 differentially expressed genes are represented as in terms of fold change. The axes are as described in (B). Yellow dot represents position of *FMR1* mRNA. Images B and C were obtained from iPathwayGuide (http://www.advaitabio.com/ipathwayguide).

In parallel, we also performed RNA-seq to identify if there were any significant transcriptional differences between our FXS hESC line and our control hESC line (Fig. 7B). A total of 1797 genes were found to be differentially expressed between untreated WT and FXS hESCs. As expected, *FMR1* expression was much lower in the FXS hESCs. Gene Ontology analysis comparing the WT and FXS hESCs datasets identified nervous system development and neurogenesis as particularly different between these two hESC lines (Suppl. Fig. 1A). Additionally, differentially expressed genes between these two hESC lines significantly map to cancer pathways (Suppl. Fig. 1B). This data is consistent with studies suggesting that FMRP regulates mRNAs involved in cancer progression and metastasis (Lucá, Averna et al. 2013, Zalfa, Panasiti et al. 2017). However, one must be cautious in interpreting these differences in expression as resulting from the *FMR1* repeat expansion or loss of FMRP as these two hESC lines are not isogenic. Of note, treatment with CGG g gRNA and dCas9-VP192 did not significantly revert FXS hESCs back towards the WT hESC transcriptome profile (data not shown).

### dCas9-VP192 activation does not increase FMRP levels in FXS hESC

In order to test whether the increase in *FMR1* transcript levels would cause a subsequent increase in FMRP, we tested the early and late FXS hESC lines with the promoter pool and CGG targeted gRNAs along with dCasVP-192. Despite a significant increase in mRNA, we did not observe a statistically significant increase in FMRP levels in either early or late passage FXS hESCs (Fig. 8A-D). Similar results were obtained with ICC measurements in these cells (data not shown). Thus, there is a dissociation at least in these cells between transcriptional reactivation and recovery of FMRP expression.

**Figure 8:**
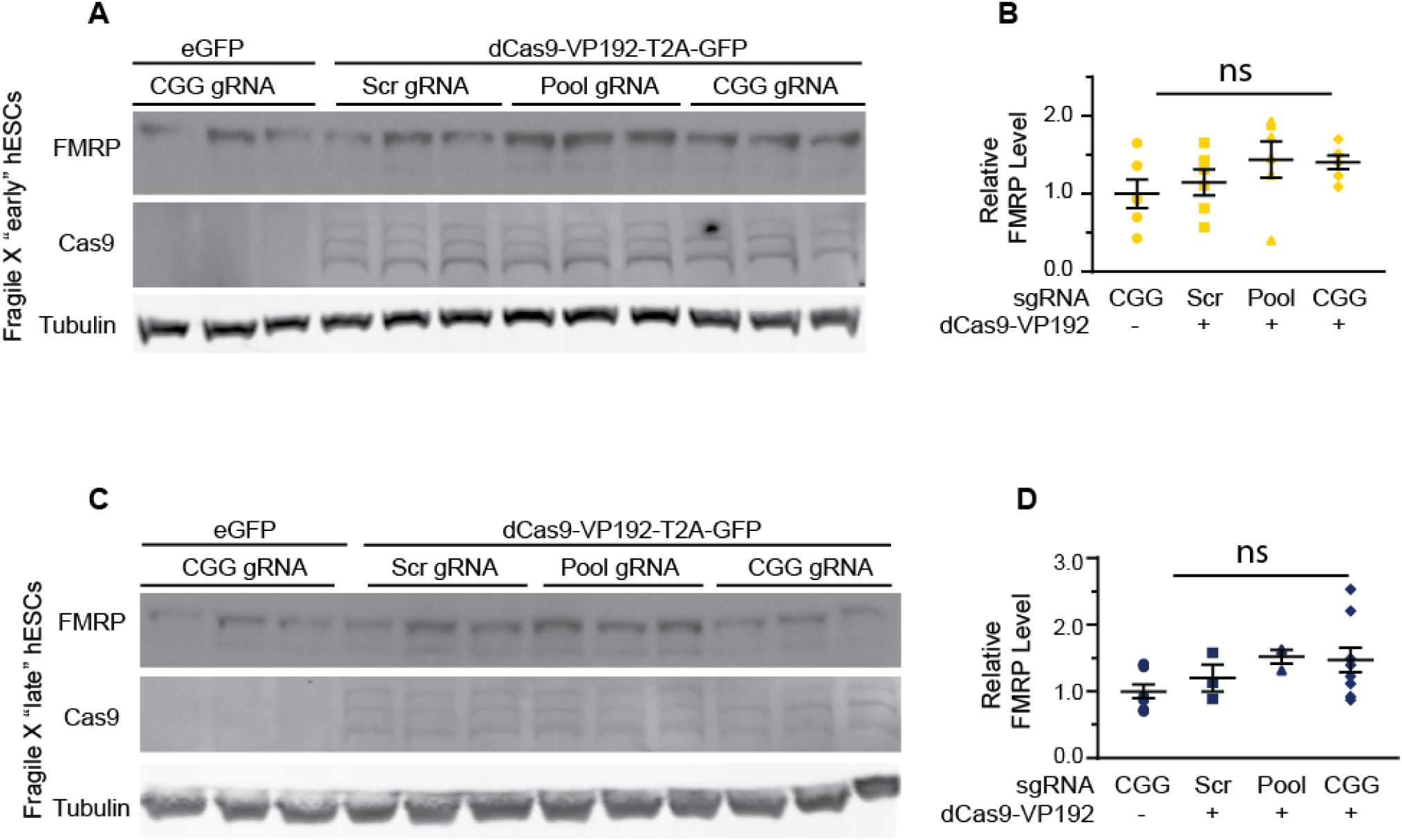
Targeted activation of *FMR1* does not significantly enhance FMRP expression in FXS hESCs. A) Western blots showing triplicate samples of early (P25-28) passage FXS hESCs transfected with control plasmid (eGFP) or with dCas9-VP192 and gRNAs immunoblotted for FMRP (by Femto ECL), Cas9 and tubulin as a loading control. B) Quantification of western blots from early passage FXS hESCs transfected as indicated. C) Western blots showing triplicate examples of late (P47-64) passage FXS hESCs transfected as in (A). D) Quantification of western blots from late passage FXS hESCs transfected as indicated.

## DISCUSSION

Fragile X Syndrome (FXS) results primarily from transcriptional silencing of the *FMR1* locus. Here, we report reactivation of *FMR1* transcription utilizing a CRISPR-dCas9 coupled transcriptional activator selectively targeted to the expanded CGG repeat. This enhanced transcription correlates with an increase in FMRP expression in both control and FXS repeat containing hEScs and occurs in both the setting of incomplete and complete transcriptional silencing and despite DNA methylation of the locus. This transcriptional reactivation is also greatest when we use a guide RNA that directly targets the CGG repeat, and this is effect is enhanced in the setting of a large CGG repeat expansion. These findings provide proof-of-concept for a CRISPR based approach to gene re-activation in FXS patient cells with the potential for translation to *in vivo* and clinical applications.

Our approach uses a nuclease deficient Cas9 to target the CGG repeats for the reactivation of the FMR1 gene. The use of a nuclease-deficient Cas9 fused to transcriptional activators or suppressors is a powerful tool for studying genome-scale events as well as specific processes (Wang, La Russa et al. 2016). Similar transcriptional activator systems have been used previously to successfully reverse disease symptoms in mouse models of Duchenne Muscular Dystrophy, highlighting the potential applicability of this system for *in vivo* treatment of disease (Long, Amoasii et al. 2016, Nelson, Hakim et al. 2016). In our hands, dCas9-VP 16 fusion constructs show the highest activation with a VP192 fusion construct along with the CGG repeated targeted gRNA. The robust activation observed using the CGG gRNA versus a promoter-targeted construct is particularly intriguing, as it suggests that the repetitive nature of the CGG guide may serve to augment its targeting strategy by providing a promoter-proximate tiling site for dCAS9-VP192 complexes. Additionally, evaluation of potential off-targets for this gRNA suggests that the presence of a large repeat element in FXS hESCs may preclude effects at other CGG repeat sites throughout the genome. Alternatively, there may be differences between hESCs and HEK293T cells in their basal transcriptomes that make them differentially sensitive to CGG repeat targeted gRNAs. This approach of directly targeting the repeats leverages the very nature of the repeat to achieve greater efficacy and specificity and has recently been used in other repeat expansion disorders to great effect (Batra, Nelles et al. 2017, Pinto, Saxena et al. 2017).

This work characterizes a new FXS hESC line, which was recently added to the NIH registry allowing for its use in United States federally–funded research. Despite a lack of FMRP, the FXS hESCs were effectively differentiated into neural rosettes and finally neurons (Fig. 4) (Eiges, Urbach et al. 2007, Telias, Segal et al. 2013)., This line exhibits a passage dependent silencing of the FMR1 gene absent any neuronal differentiation (Fig 4). This is consistent with some published findings suggesting selection against expression of large expanded CGG repeat containing RNAs and (potentially) RAN translation products (Brykczynska, Pecho-Vrieseling et al. 2016, Zhou, Kumari et al. 2016). However, it disagrees with work in the first characterized and widely used hESC FXS line that exhibits a neuronal differentiation-dependent silencing that appears dependent on an RNA induced transcriptional silencing mechanism (Eiges, Urbach et al. 2007, Colak, Zaninovic et al. 2014). Our work does not delineate between these two possibilities, but future studies over longer time courses using stable transfection systems will be needed to determine both the sustainability of the enhanced transcription observed and the impact of enhanced production of large CGG repeat RNAs on cell viability and differentiation.

Despite retaining some transcriptional activity in the early passage state, this new FXS hESC line exhibits a methylated FMR1 locus. This finding is surprising, as previous work suggested that this methylation typically occurs coincident with or after transcriptional silencing. We explored whether this methylation might instead be by hydroxymethylation, which is not transcriptionally silencing (Brasa, Mueller et al. 2016, Esanov, Andrade et al. 2016). However, our preliminary studies demonstrate no evidence of hydroxymethylation in these FXS cells (data not shown). Future work will be needed to better characterize the epigenetic and methylation state of this locus in these early and late passage hESCs to better delineate the time-course of transcriptional silencing.

This study is complementary to a series of recent papers utilizing the CRISPR-Cas9 system to reactivate transcription from the *FMR1* locus in FXS. Two studies took a more direct approach of cutting out the repeat with the Cas9 nuclease and both achieved correction ofthe transcriptional silencing and a reactivation of FMRP expression (Park, Sung et al. 2016, Xie, Gong et al. 2016). More recently, a third study used a strategy more akin to our approach, targeting gRNAs to the CGG repeat and coupling that with a dCas9 fused to the active domain of the TET DNA demethylase (Liu, Wu et al., Liu, Wu et al.). Using this approach in an iPSC line with ~500 CGG repeats, they were able to achieve both transcriptional reactivation of FMR1 as well as at least partial recovery of FMRP expression (Liu, Wu et al.). As with our work, it is intriguing in all of these studies that reactivation of FMR1 transcription can occur even at a fully methylated and transcriptionally silenced locus observed in the late passage FXS hESCs (Fig. 4B). This suggests that methylation and heterochromatization of the locus do not preclude access of the gRNAs to the repeat sequence. Our study adds the additional element that even targeting a transcription factor to the repeat, which does not directly target any of the epigenetic alterations present at the locus in FXS, is sufficient to reactivate the gene. Taken together, these findings imply that the silenced CGG repeat expanded FMR1 locus may be more dynamic than previously thought-at least in the setting of hESCs where such boundaries may be more permissive to epigenetic change. Moreover, these results imply that FMR1 transcriptional reactivation can be achieved through multiple potentially complementary approaches.

While the dCas9-VP192 activation system in control HEK293T cells elicited relatively equivalent effects on both *FMR1* transcription and FMRP production, in hESCs with pathologic repeat expansions the impact of transcriptional upregulation on FMRP expression was significantly blunted (Fig. 7). This effect is consistent with large body of research demonstrating that expanded repeats acts as an impedance to ribosomal scanning and downstream initiation of FMRP translation(Feng, Zhang et al. 1995, Chen, Tassone et al. 2003, Khateb, Weisman-Shomer et al. 2007, Iliff, Renoux et al. 2013). How large of a factor such a translational blockade might play in any transcriptional re-activation strategy is unclear. Very large unmethylated repeats that are efficiently transcribed can still produce a FXS phenotype, although some cases of methylation mosaicism and repeat length mosaicism have only modest or no clear clinical symptoms(Burman, Popovich et al. 1999, Tassone, Hagerman et al. 2000). It may be that the underlying repeat size is the critical determinant. Most cases of fully unmethylated full mutation patients described to date have repeat sizes that are less than 400 CGGs (Pietrobono, Tabolacci et al. 2005, Tabolacci and Chiurazzi 2013). In cases of methylation mosaicism, somatic instability complicates data interpretation, meaning that effects on FMRP production may be cell specific (Jiraanont, Kumar et al. 2017). If large repeats preclude recovery of FMRP expression in FXS patients with very large repeats, then approaches specifically targeting this translational blockade will be needed to achieve reactivation. However, given that less CGG DNA methylation, more *FMR1* mRNA transcription and more FMRP production in even a subset of cells in FXS patients all correlate with better clinical outcomes and differential responses to pharmacological agents, even modest successes targeting these proximal events in pathogenesis may elicit meaningful effects on clinical phenotypes. Thus, this proof-of principal study provides hope that such approaches will eventually lead to effective therapeutics in patients with Fragile X syndrome.

## DATA AVAILABILITY

The datasets generated for this study will be posted to the GEO database upon formal publication.

## AUTHOR CONTRIBUTIONS

PT and JMH conceived the project. JMH conducted all of the experiments in HEK293T cells and all initial experiments in hESC cells. GS validated hESC experiments did NPC and neuronal work and performed the RNA-seq experiments. GS wrote the manuscript with significant input from PT and JMH. GDS and AMR derived the UM139-2 hESC line. CR initially characterized the UM 139-2 hESC line. JP provided assistance with stem cell work. All authors edited the manuscript.

## ACKNOWLEDGEMENT

The authors thank Eric Wang and Lance Denes for technical support and helpful discussions. We thank the University of Michigan Neurology Stem Cell core and MStem Cell lab for assistance with stem cell culture work specifically Sandra Mojica-Perez, Jun Ding and Laura Keller. We thank the UM Bioinformatics core for library prep and assistance with RNA-seq data.

---

**Figure S1:**
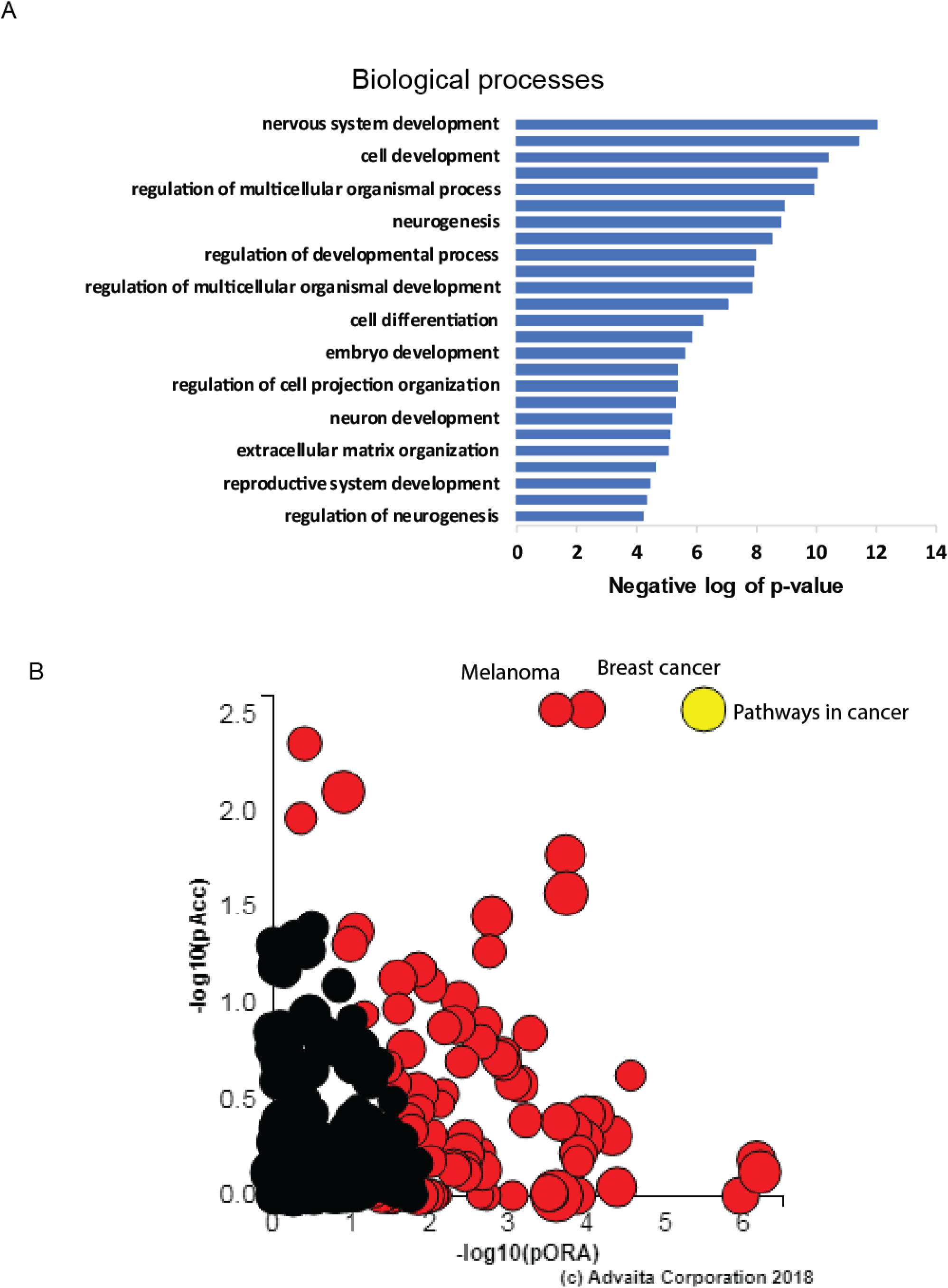
Gene Ontology analysis and pathways for WT versus late FXS hESCs. A) Graph depicting top Gene Ontology (GO) terms for the RNA-seq data comparing WT and FXS hESCs. X-axis shows the GO terms and Y-axis shows the negative log (base 10) of the p-values for each category. B) All pathways from the WT and FXS hESC RNA-seq analysis are plotted in terms of the two types of evidence computed by iPathwayGuide using Impact Analysis (Tarca, Draghici et al. 2009): over-representation on the x-axis (pORA-Over Representation Analysis) and the total pathway accumulation on the y-axis (pAcc-Accumulated perturbation of the pathway). Each pathway is represented by a single dot, with significant pathways shown in red, non-significant in black. Both p-values are shown in terms of their negative log (base 10) values. Yellow dot represents cancer pathways. The adjacent red dots show pathways for melanoma and Breast cancer. Figure obtained from iPathwayGuide (http://www.advaitabio.com/ipathwayguide).

